# PHOSPHATE OVERACCUMULATOR 2 (PHO2) is a negative regulator of arbuscular mycorrhizal symbiosis

**DOI:** 10.1101/2025.09.03.673468

**Authors:** Saskia Birch, Sophie Perryman, Evan E. Ellison, Nina Foreman, Nikoline Mekjan, Alex Williams, Miles Bate-Weldon, Thomas Ralfs, Boas Pucker, Michelle Whiting, Matthew S. Hope, Chongjing Xia, Emma Wallington, Katie Field, Jeongmin Choi

## Abstract

Arbuscular mycorrhizal (AM) symbiosis is an ancient relationship formed between most plants and Glomeromycotina fungi, typically in response to phosphate (Pi) limitation in soils. By hosting these fungi in their roots, plants extend their access to essential mineral nutrients and water beyond the rhizosphere, while providing the fungus with carbon in return. This mutualistic symbiosis presents a promising tool for enhancing sustainability in agriculture, as it not only supports plant nutrition but also immunity and wider soil health. However, achieving high crop yields currently relies on supplementing plants with excess Pi, which suppresses AM symbiosis. We found that this suppression is mediated by a key negative regulator of the Pi starvation response (PSR) in rice (*Oryza sativa*), Phosphate overaccumulator 2 (PHO2). *PHO2* encodes an E2 ubiquitin-conjugating enzyme which targets various proteins involved in the PSR in Pi-sufficient conditions for protein degradation. Here we report that *pho2* mutants of rice and *Nicotiana benthamiana* retained high AM fungal colonisation even in high Pi conditions. Our transcriptomic analysis of uninoculated rice roots revealed that *pho2* mutants are less sensitive to Pi treatment and retain susceptibility to AM symbiosis by maintaining expression of a core set of AM-related genes gating the early stage AM fungal entry, such as genes involved in strigolactone biosynthesis, LysM-containing plant receptors for fungal molecules, and components of the common symbiosis signalling pathway (CSSP). Furthermore, isotope tracing using ^33^P and phosphate transporter (*PHT1*) gene expression patterns collectively suggest enhanced direct and symbiotic Pi overaccumulation in *pho2* mutant leaves. Together, our data reveal a new role for PHO2, as a negative regulator of AM colonisation and symbiotic Pi accumulation in shoots.

## Introduction

Phosphorus (P) is an essential nutrient for plant growth and survival. Plants acquire P as inorganic phosphate (Pi), which is found in low abundance in most soils. To increase access to this scarce nutrient, around 80 % of land plants engage with *Glomeromycotina* fungi in arbuscular mycorrhizal (AM) symbiosis ^1^. Arbuscular mycorrhizal (AM) fungi establish transient tree-like ‘arbuscules’ to facilitate nutrient exchange within plant cortical root cells ^2^. Their extraradical hyphal network extends beyond the rhizosphere to supply their host with mineral nutrients in exchange for plant-derived carbohydrates and fatty acids required to complete their life cycles ^3, 4, 5^. Critically, AM symbiosis can contribute up to 70-99 % of a plant’s Pi nutrition ^6, 7, 8^.

Plants experiencing Pi depletion engage AM fungi by secreting phytohormones called strigolactones (SLs) and other compounds from their roots. This induces fungal germination and hyphal growth towards plant roots ^9, 10, 11, 12^. AM fungi reciprocate this pre-symbiotic contact by releasing chitin-derived molecules known as myc factors ^13, 14^. These are perceived by lysine motif (LysM) –containing plant receptors at the root, which trigger a downstream signalling cascade culminating in transcriptional reprogramming to accommodate symbiosis ^15, 16^. AM fungi form infection structures known as hyphopodia on the surface of the plant roots before entering and traversing the root cortex. This is facilitated by the common symbiosis signalling pathway (CSSP) which similarly mediates nitrogen-fixing root nodule symbiosis by rhizobacteria ^17^. Within the inner cortical cells, AM fungi form highly branched arbuscules which act as sites of nutrient exchange^18^.

To ensure the mutualistic nature of mycorrhizal nutrient exchange, plants suppress engagement with AM fungi in high Pi conditions, instead favouring direct nutrient uptake by Pi transporters on the root^19^. Recent studies have started to uncover the molecular underpinnings of this Pi-dependent regulation of AM symbiosis ^20, 21, 22^. In both rice (*Oryza sativa*) and *Lotus japonicus,* a highly conserved Pi starvation response (PSR) pathway has been shown to play a central role in promoting AM symbiosis in low Pi conditions ^20, 21^.

The PSR centres around MYB transcription factors (TFs), PHOSPHATE STARVATION RESPONSE (PHRs), which regulate a large proportion of plants’ adaptations to Pi-deficiency ^23, 24, 25, 26^. PHRs bind to a conserved PHR1 binding site (P1BS) in the promoter regions of their target genes to activate their transcription ^24, 25^. This promotes expression of various gene products involved in Pi uptake and remobilisation, including the Phosphate transporter 1 (PHT1) family of high-affinity Pi transporters, purple acid phosphatases, and regulatory non-coding RNAs ^23, 24, 27, 28, 29, 30, 31, 32^. PHR TFs in rice and *L. japonicus* also upregulate genes involved in establishing AM symbiosis in low Pi conditions, even in the absence of AM fungi ^20, 21, 22^. These genes regulate various stages of symbiotic establishment, including SL biosynthesis, the CSSP, and nutrient transport. The *phr2* mutant in rice displays substantially reduced AM colonisation in Pi-depleted plants ^20, 21^. In contrast, overexpressing *PHR2* in rice and *L. japonicus* partially restores colonisation in Pi-sufficient conditions which suppress colonisation of wild-type (WT) plants ^20, 21^.

In high Pi conditions, SPX (named after its homologs, *SYG1/Pho81/XPR1*) proteins suppress PHR activity ^30, 33, 34, 35, 36^. SPXs bind to inositol pyrophosphates (InsPs), which accumulate with increased levels of intracellular Pi ^37^. In complex with InsPs, SPXs bind to PHRs via a coiled-coil domain in the PHR protein ^30, 33, 35, 36, 38, 39^. This blocks PHRs from homodimerization, entry to the nucleus and DNA binding ^30, 34, 35, 36, 38, 40^. Rice *spx1/2/3/5* and quadruple mutants and tomato *spx1* mutants exhibit increased AM colonisation of roots compared to WT; conversely, overexpression of *SPX* reduces colonisation, indicating that SPXs also suppress AM colonisation ^21, 41^.

Another key negative regulator of the PSR in Pi-sufficient conditions is Phosphate overaccumulator 2 (PHO2), an E2 ubiquitin-conjugating enzyme ^23, 42, 43, 44^. Its targets include high-affinity Pi transporters, PHT1s; Phosphate 1 (PHO1), a transporter responsible for loading Pi into the xylem; and PHOSPHATE TRANSPORTER TRAFFIC FACILITATOR1 (PHF1) which mediates the shuttling of PHT1 Pi-transporters to the plasma membrane ^45, 46, 47, 48, 49^ and has also been implicated in a wide range of other PSR-related activities^44^. The overaccumulation of PHO2 target proteins lead to a typical *pho2* mutant phenotype, culminating in hyperaccumulation of Pi in its shoots and impaired remobilisation of Pi from old to new leaves ^23, 42, 43, 44^. Consequently, rice *pho2* mutants are shorter than WT plants and display leaf tip necrosis, which is a symptom of Pi toxicity ^43, 44^.

In Pi-depleted conditions, *PHO2* is suppressed by the miR399 family of microRNAs ^23, 42, 44, 46, 47^, which are activated by PHRs ^23, 28^. These miRNAs suppress *PHO2* transcriptionally by binding to its mRNA and targeting it for degradation ^23, 28, 42^. This is counteracted by the activity of a PHR-activated long non-coding RNA, *INDUCED BY PHOSPHATE STARVATION 1* (*IPS1*) which mimics the miR399 target sequence with a 3 nucleotide bulge at the cleavage site, causing sequestration ^4, 27, 31^. These regulators fine-tune activity of the PSR in response to a dynamic range of Pi levels. However, the role of PHO2 in regulating AM symbiosis in remains unknown.

In this study, we found that PHO2 functions as a suppressor of AM colonisation in rice and *Nicotiana benthamiana* in high Pi conditions. By using a combination of transcriptomics and isotope tracing techniques in rice, we revealed a new role of PHO2 in regulating early establishment of AM colonisation and symbiotic Pi accumulation in shoots.

## Results

### PHO2 suppresses AM colonisation in high Pi conditions

The rice genome contains a single copy of the *PHO2* gene (**Fig.S1**) ^23^. To uncover the role of PHO2 in AM symbiosis in high Pi conditions, we obtained two independent Tos17 transposon lines, *pho2-1* (NF2586) and *pho2-2* (NE9017), in the Nipponbare cultivar (Rice Genome Resource Centre, Japan; **Fig.S2a**). In both lines, the transposons were inserted in the beginning of the first exon, expected to impair PHO2 function. Indeed, quantitative reverse transcription (qRT-)PCR showed that expression of *PHO2* is significantly lower in *pho2-2* mutants (**Fig.S2b**), while *pho2-1* and *pho2-2* mutants accumulated higher levels of Pi in the shoots (**Fig.S2c**), confirming the previously reported phenotype of *pho2* mutants in rice ^43, 44^. In WT, root Pi concentration also increased with increasing Pi treatment; however, *pho2* mutants retained the same root Pi concentration regardless of Pi treatment (**Fig.S2d**). This provides further evidence that *pho2* mutants prioritise Pi accumulation in their shoots, although the difference in root Pi concentration between genotypes is too minor to see a statistically significant genotype effect in either Pi treatment.

Subsequently, to determine the degree to which AM fungal colonisation was affected in WT and *pho2* mutant plants, we inoculated seedlings with 300 spores of the AM fungal strain, *Rhizophagus irregularis,* and grew them in three increasing concentrations of Pi (25 µM, 250 µM and 1500 µM). In low Pi (25 µM) conditions, all genotypes successfully established AM colonisation (**Fig.1a**) although *pho2* mutants had higher arbuscule abundance (**Fig.1a, b**). At 250 µM Pi, both total root length colonisation and arbuscule presence were severely decreased in WT. Remarkably, at 250 µM Pi both *pho2* mutants maintained high total and arbuscule colonisation levels (80-100 %) (**Fig.1a**). At unusually high Pi levels (1500 µM) AM symbiosis decreased in the *pho2* mutants, although *pho2-2* was not as severely affected as the *pho2-1* mutant, in which colonisation decreased to WT-like levels. The differing severity of *pho2* phenotype between mutant alleles was correlated with endogenous *PHO2* expression level, although both alleles showed similar alterations in phosphate starvation marker gene expression (i.e. *IPS1*, **Fig.S2b**). This Pi dose-dependent *pho2* phenotype indicates a Pi threshold within which PHO2 functions as a suppressor of AM colonisation.

**Figure 1.**
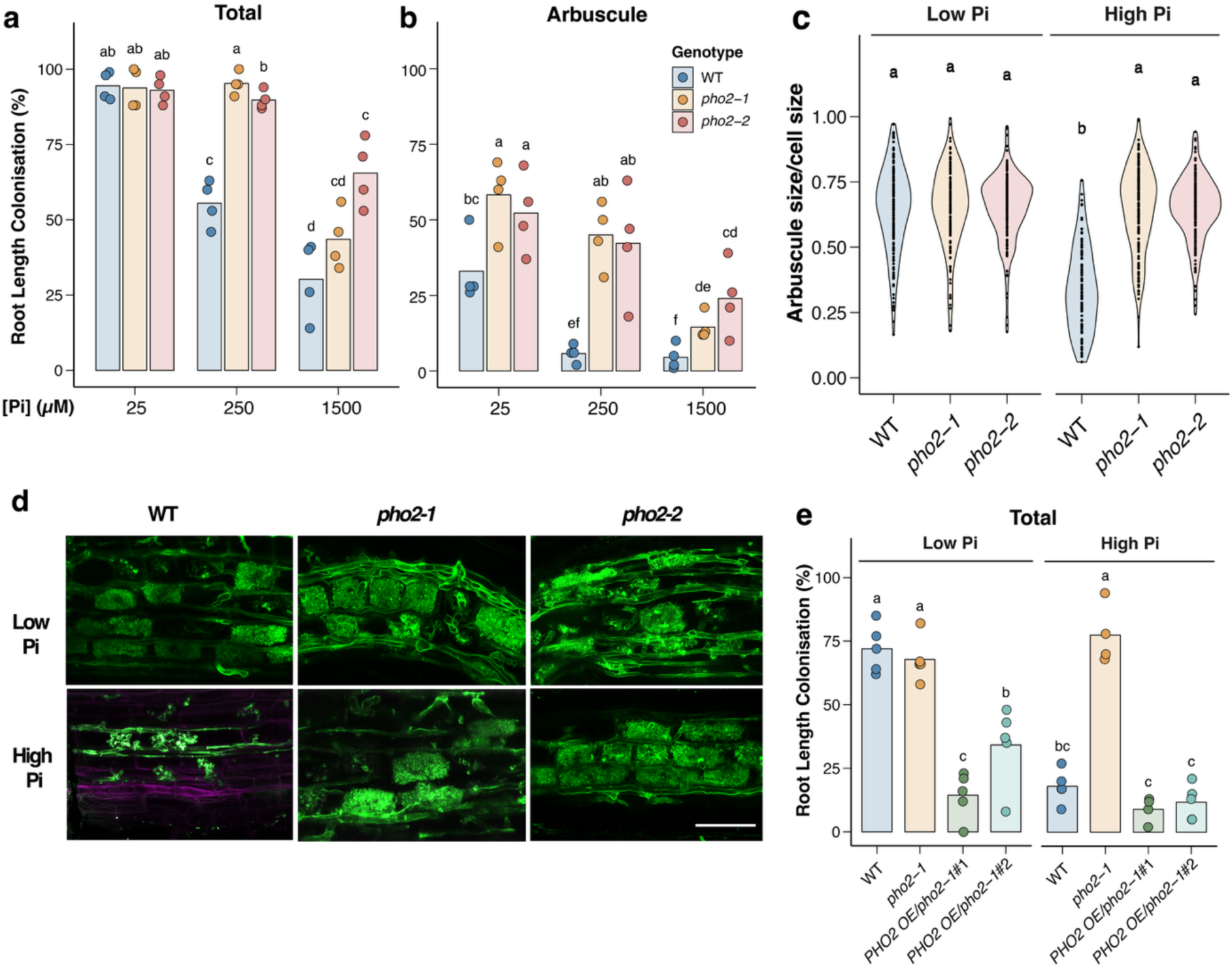
AM colonisation in rice *pho2* mutants. **a)** Root length colonisation of total and **b)** arbuscules. **c)** Arbuscule size relative to cell size **d)** confocal images of arbuscules stained with WGA-Alexa488 and propidium iodide, in wildtype (WT) and 2 alleles of *pho2* mutant rice plants. Plants were grown in cones for 6 weeks post inoculation (wpi) with 300 spores of *Rhizophagus irregularis* and received half-strength Hoagland’s solution containing low and high Pi indicate Pi concentrations of 25 or 250 µM, respectively. **e)** Total root length colonisation of two *PHO2* overexpression lines expressing *PHO2* CDS under the maize Ubiquitin promoter in the *pho2-1* background. Bars indicate the mean value for each group and points show individual plant as a biological replicate. For statistical analysis, Kruskal-Wallis tests were performed (**(a)** p-value = 0.00261, n = 4; **b)** p-value = 0.000368, n = 4; **c)** p-value = 0, n = 150-200 (n = 105, in high Pi, WT); **e)** p-value = 0.000118, n = 4 or 5). Letters indicate significantly different groups (p < 0.05) based on post-hoc pairwise comparison using the Agricolae package in RStudio. **d)** Scale bar indicates 50 µm.

To identify the stages of AM colonisation which are affected by PHO2-mediated suppression, we performed a time-course experiment in low Pi (LP; 25 µM) and high Pi (HP; 250 µM) conditions. Interestingly, AM colonisation was higher in the *pho2* mutants compared to WT at an early time-point (4 weeks post inoculation, wpi) in LP conditions (**Fig.S3**), indicating that PHO2 may suppress early establishment of AM symbiosis. Furthermore, while WT plants reached a peak colonisation at 6 wpi in LP conditions, they never achieved the same level of root colonisation as the *pho2* mutants (**Fig.S3**). This alludes to a regulatory role of PHO2 in LP conditions as well as in HP. In HP conditions, AM colonisation levels of the *pho2* mutants displayed striking contrast to WT: AM colonisation in *pho2* mutants increased consistently up until 6wpi, reaching around 75 %, whereas WT plants were less than 20 % colonised throughout the time-course (**Fig.S3**). This highlights the indispensable role of PHO2 in suppression of AM colonisation in HP conditions.

The highly branched nature of arbuscules provides a large surface area for nutrient exchange. It has been reported that mutants of another PSR component in rice, *phr2*, form smaller collapsed arbuscules^21^ which are less able to perform symbiotic Pi uptake. To evaluate whether arbuscules in *pho2* in HP are similarly dysfunctional, we examined the arbuscule shape and size in *pho2* mutants at 6wpi. In LP conditions there were no significant differences in arbuscule size and shape between WT and *pho2* mutants (**Fig.1c, d**). However, in HP conditions, arbuscules were larger in the *pho2* mutants than in WT (**Fig.1c**) and we observed more collapsed arbuscules in WT (**Fig.1d**). These findings suggest that signalling downstream of PHO2 may suppress arbuscule formation or promote arbuscule collapse and turnover.

To ensure that PHO2 is responsible for the observed AM colonisation phenotype, we generated two *PHO2* over-expression (OE) lines, *PHO2-*OE1 and *PHO2-*OE2, expressing the *PHO2* CDS under the control of a maize ubiquitin promotor in the *pho2-1* mutant background. These plants were inoculated with *R. irregularis* and grown in LP and HP conditions to assess their root colonisation levels. The expression levels of *PHO2* in roots of the *PHO2*-OE1 and *PHO2*-OE2 were ∼ 30-fold and 6.5-fold higher than WT, respectively (**Fig.S4a**). In HP conditions, the total colonisation level of both *PHO2* OE lines were successfully restored to WT levels (< 20 % root length colonisation), while in *pho2-1* mutants colonisation level remained elevated. Interestingly, overexpressing *PHO2* suppressed AM colonisation levels even in the permissive LP conditions, possibly due to the constitutive action of the maize ubiquitin promoter (**Fig.1e**). The root length colonisation of arbuscules followed a similar pattern to the total colonisation level (**Fig.S4b**). Collectively, this genetics data confirms that PHO2 is indeed responsible for the observed suppression of AM colonisation in HP conditions.

### AM colonisation does not promote shoot growth in *pho2* mutants

AM colonisation can promote growth and nutrient uptake in WT rice ^50^. To assess whether this occurs in *pho2* mutants, plants were inoculated with 600 *R. irregularis* spores in 400 mL pots for 8 weeks under LP (25 µM) and HP (250 µM) conditions and compared with uninoculated controls. As reported previously ^43, 44^, the *pho2* mutant plants were smaller than WT in both phosphate conditions. WT plants exhibited increased shoot fresh weight upon AM colonisation, however, this growth promotion effect was not observed in HP conditions where AM colonisation was inhibited (**Fig.2a**). In contrast, the *pho2* mutant did not exhibit any growth promotion in either LP or HP conditions (**Fig.2a**), despite maintaining high root colonisation and fully developed arbuscules in both LP and HP conditions (**Fig.S5a, b**).

**Figure 2.**
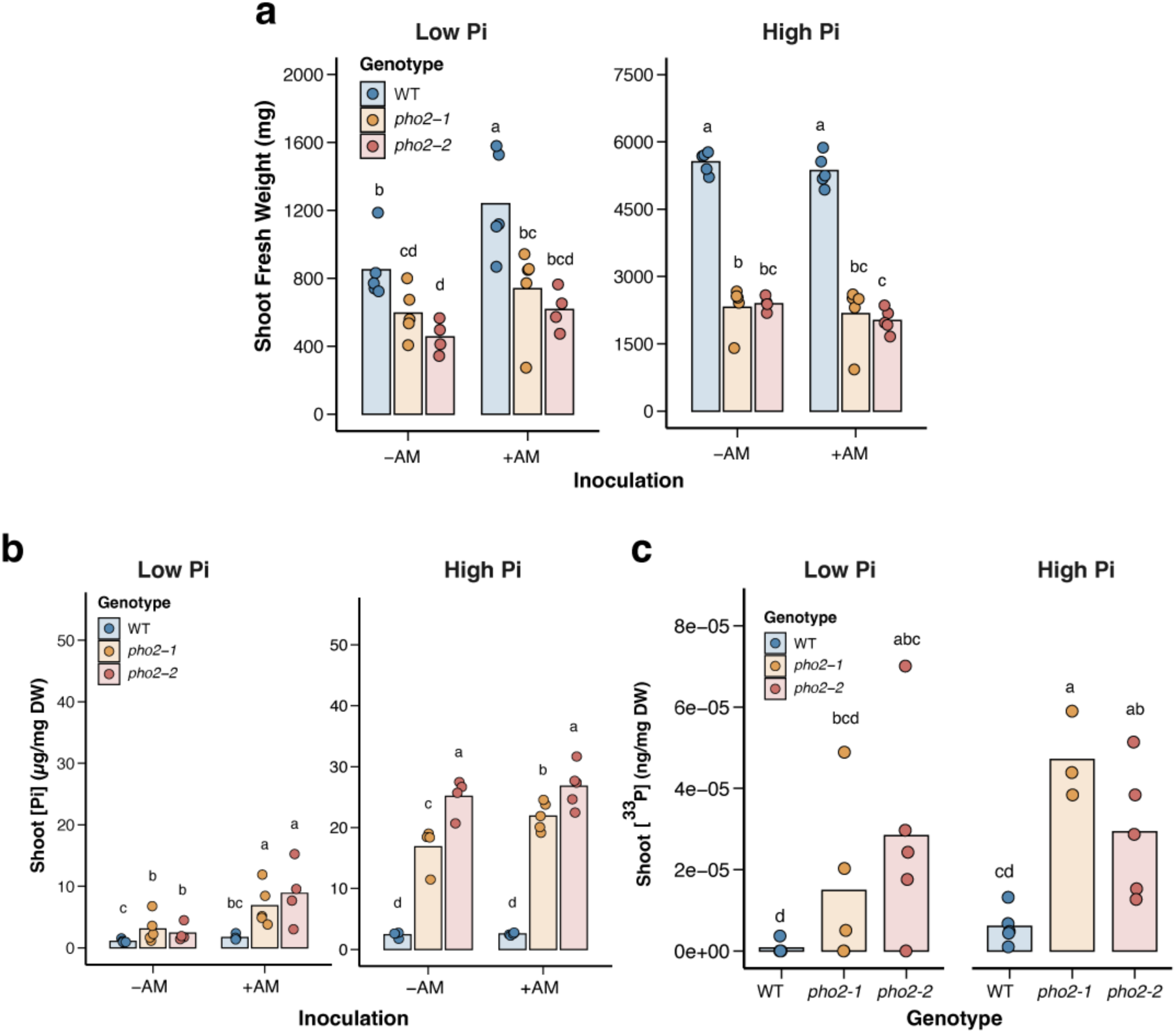
Mycorrhizal growth promotion and phosphate levels in shoots. **(a)** Shoot fresh weight and (**b)** shoot phosphate (Pi) concentrations of uninoculated (-AM) and inoculated (+AM) WT and *pho2* mutant rice plants (*pho2-1* and *pho2-2).* Plants were grown in pots for 8 weeks following inoculation with 600 spores of *Rhizophagus irregularis and* fertilised with half-strength Hoagland’s solution containing 25 (low Pi) or 250 µM (high Pi) phosphate. (**c)** Concentration of ^33^P obtained via mycorrhizal hyphae in shoots of plants grown in 1 L pots with low and high Pi. At 8 weeks post-inoculation, ^33^P-orthophosphate was administered to an AM-accessible core in each pot, and plants were harvested 2 weeks later. Bars signify the mean value for each group and points show individual samples. Kruskal-Wallis tests were performed (**a)** low Pi: p-value = 0.00390; high Pi: p-value = 0.000741 and n = 4 or 5; **b)** low Pi: p-value=0.00133; high Pi: p-value = 0.000342 and n = 4 or 5; **c)** p-value = 0.00987, n = 3, 4 or 5) and post-hoc pairwise comparisons using RStudio’s Agricolae package generated letter labels to indicate significantly different groups (p < 0.05).

Interestingly, Pi concentrations in WT and *pho2* mutants did not correlate with a visible growth promotion (**Fig.2b**). In both LP and HP conditions, WT plants maintained constant Pi concentrations in their shoot with or without AM inoculation (**Fig.2b**). Contrastingly, the *pho2* mutants had a higher shoot Pi concentration when inoculated with *R. irregularis* compared to uninoculated controls in LP conditions (**Fig.2b**). In addition, in HP conditions, only *pho2-1* showed significantly increased shoot Pi concentration with AM-inoculation. This provides the first evidence that PHO2 may play a role in Pi transport from roots to shoots and maintaining optimal shoot phosphate homeostasis during AM symbiosis.

### Symbiotic Pi transfer is maintained in HP conditions

To evaluate phosphate transfer activity of *pho2* arbuscules in HP conditions, we traced movement of ^33^P-labelled orthophosphate from AM fungi to host plant tissues by introducing the isotope to a 35 µm nylon mesh-covered core that exclusively permits access by fungal hyphae but not plant roots ^51^. To control for plant assimilation of the isotope via passive diffusion and/or alternative microbial P-cycling processes, the isotope-containing core in half of the pots was rotated to sever AM fungal connections between the host plant and the core contents. Plants were grown for 8 weeks to fully establish AM colonisation (**Fig.S6a, b**) and then allowed to grow for a further 2 weeks following the addition of 1MBq ^33^P-orthophosphate to the hyphal core. Total phosphorus (P) and AM fungal-acquired ^33^P content of WT and *pho2* mutant plants were quantified in the plant shoot tissues.

As previously observed, the *pho2* mutants were significantly smaller than WT plants in both LP and HP conditions and their biomass was only slightly increased by treatment with HP (**Fig.S6c**). Similarly to Pi concentration, the concentration of total P (both directly and symbiotically-acquired) was higher in the shoots of *pho2* mutant plants compared to WT in both LP and HP treatments (**Fig.S6d**). Symbiotic delivery of ^33^P was maintained in WT and *pho2* mutant plants in HP conditions, even though AM colonisation was reduced in WT in HP (**Fig.2c**). In LP conditions, *pho2* mutants had higher ^33^P concentration in their shoots, indicating greater mycorrhizal Pi uptake per unit biomass despite comparable colonisation levels (**Fig.S6a**). ^33^P concentration showed similar elevation in *pho2* mutant shoots compared to WT in HP conditions, likely due to their higher AM colonisation. This reinforces our findings regarding Pi concentrations and provides further evidence for our hypothesis that PHO2 regulates symbiotic-acquired Pi contents in the shoots.

### PHO2 suppresses genes involved in pre-symbiotic establishment and the common symbiosis signalling pathway

The *pho2* mutants of both *Arabidopsis* and rice have been found to have substantial changes in gene expression compared to WT plants ^23, 42, 44^. This suggests that the downstream effects of PHO2 lead to transcriptional reprogramming. However, the *pho2* mutant transcriptome is yet to be characterised in the presence and absence of AM symbiosis. To explore how PHO2 impacts the expression of AM-related genes, we performed transcriptional analysis of WT and *pho2-1* (henceforth, *pho2*) roots grown in LP (25 µM) and HP (250 µM) conditions from the Pi gradient experiment (**Fig.1a**).

The number of differentially expressed genes (DEGs) showed dramatically altered gene expression in *pho2* compared to WT plants, especially in HP conditions and when inoculated with AM fungi (**Fig.3a**); this finding is reflected in the elevated AM-colonisation levels in the *pho2* mutant in HP conditions (**Fig.1a**). Consistent with these findings, the *pho2* mutant maintains quite a significant number of DEGs in response to mycorrhizal inoculation in HP compared to the dampened transcriptional response in WT (**Fig.3a**). Crucially, the number of DEGs in HP compared to LP reveals that *pho2* mutants have 93.2 % and 96.2 % fewer up-and down-regulated DEGs than WT in uninoculated (mock) conditions, illustrating the insensitivity of *pho2* mutants to external Pi treatment.

**Figure 3.**
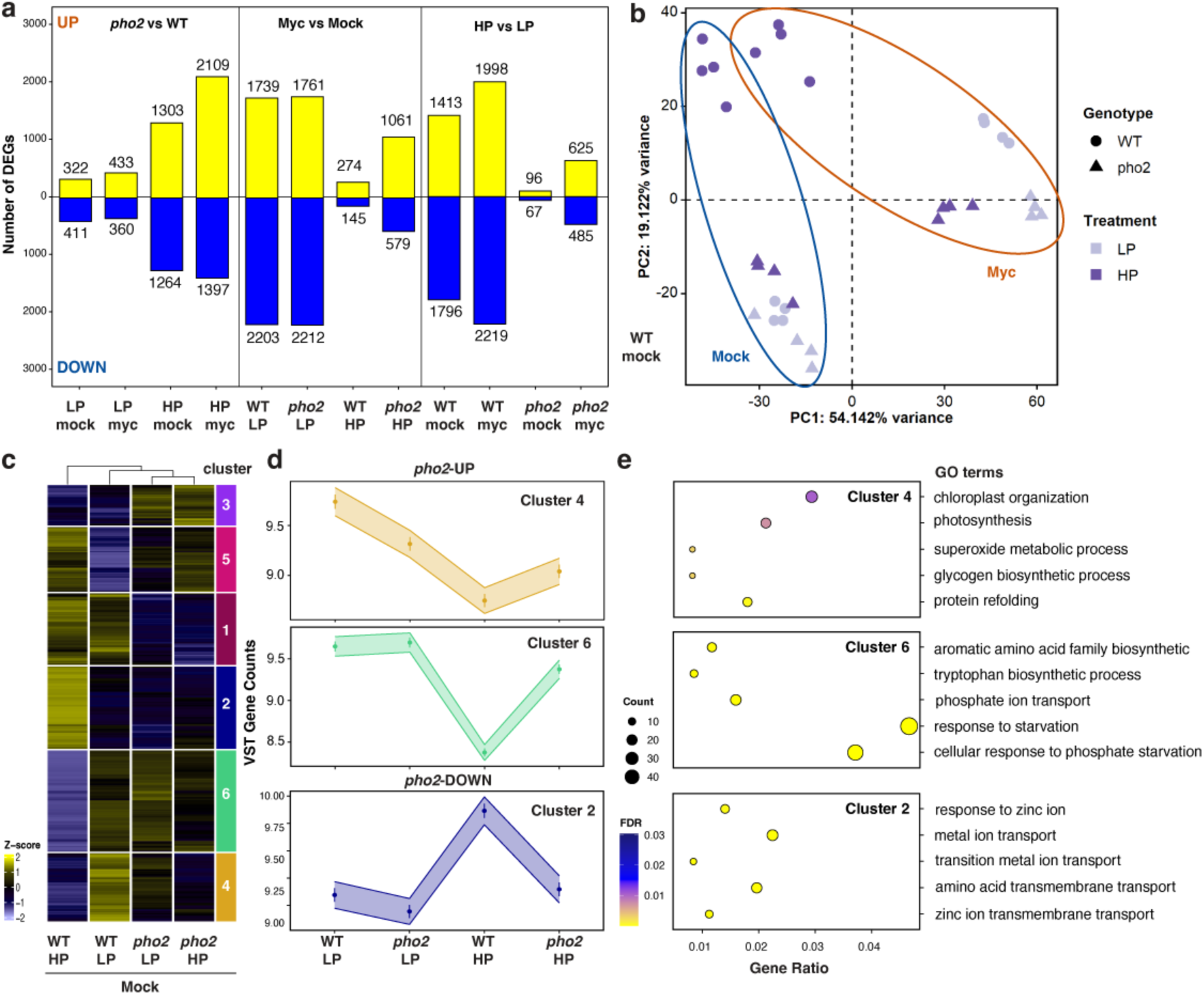
Transcriptomic analyses of *pho2* mutant roots. Plants were grown for 6 weeks and fertilised with half-strength Hoagland’s solution containing 25 (LP) or 250 (HP) µM phosphate. Four biological replicates were used in each group for RNA-seq, and each replicate is from a single plant**. (a)** The number of differentially expressed genes (DEGs) based on the indicated pairwise comparisons (FDR ≤ 0.05 and log_2_FoldChange ≥ log_2_(1.5)). (**b)** Principal component analysis (PCA). (**c)** Hierarchical clustering of DEGs in uninoculated samples using k-means clustering of samples. Yellow and blue indicates up- and down-regulated genes across genotype per gene. (**d)** Representative expression patterns of Clusters 4, 6 and 2 using median VST gene counts of all genes in each cluster. The shaded area shows ± standard deviation from the median. The x-axis denotes the genotype and phosphate treatment (**e)** Top 5 categories of Gene Ontology enrichment (GO) of clusters 4, 6 and 2.

Furthermore, a principle component analysis (PCA) was conducted to identify related transcriptional signatures. Notably, uninoculated (mock) and AM-inoculated (myc) samples form distinct groups, except for in WT in HP conditions where mock and myc samples are grouped together to form their own defined cluster. Interestingly, the *pho2* mutant treated with HP clustered with all LP samples, potentially suggesting that the reduced sensitivity of the *pho2* mutant is due to constitutively active phosphate starvation status in *pho2* regardless of Pi treatment.

To narrow down PHO2-regulated genes involved in AM colonisation, we performed hierarchical clustering to identify genes displaying similar patterns of expression which correspond to AM colonisation. Intrigued by elevated early infection levels from our time-course experiment (**Fig.S3**), we focused on the transcriptome of uninoculated roots to examine host permissibility to AM fungi prior to physical contact. Cluster analysis successfully grouped related root samples with similar biological characteristics. For example, *pho2* in LP and HP clusters closely with WT samples grown in LP (**Fig.3c**). Among six clusters, we designated clusters 4 and 6 as *pho2-*UP; these genes are suppressed in WT in HP conditions and show higher transcript levels in *pho2* compared to WT in HP conditions, positively correlating with patterns of AM colonisation **(Fig.3d**). In contrast, cluster 2 was assigned as *pho2*-DOWN, negatively correlating with AM colonisation patterns. Crucially, gene ontology (GO) enrichment showed that cluster 6, in particular, was enriched with genes involved in plant responses to phosphate starvation (**Fig.3e**); indeed, *pho2-*UP genes account for 47.3 % (850/1796) of genes that are phosphate starvation-induced (PSI) in WT (PSI-UP; **Fig.4a**). Upregulation of PSR in *pho2* was corroborated by qRT-PCR of the PSI marker gene, *IPS1* (**Fig.S2b**).

**Figure 4.**
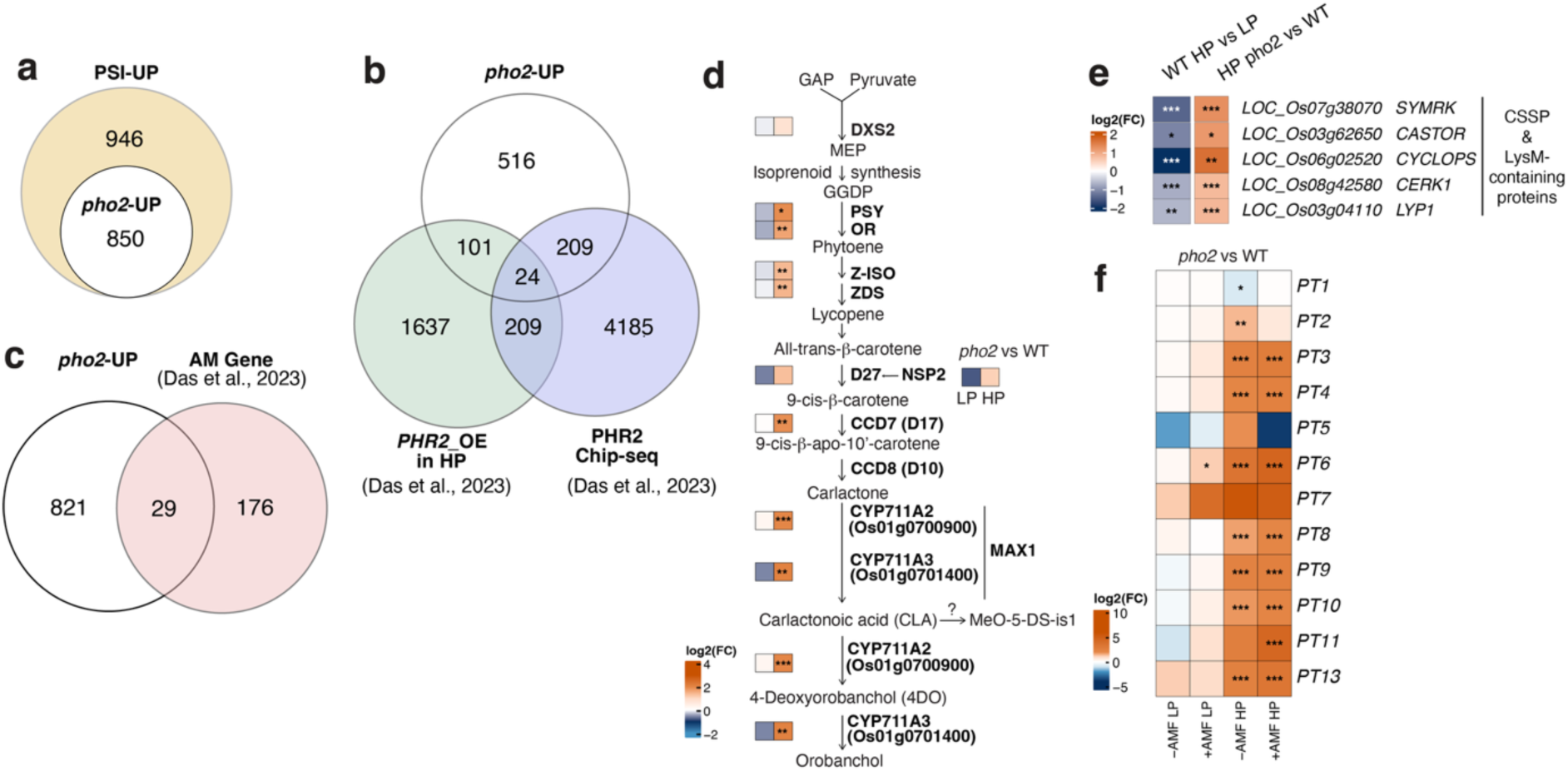
PHO2 regulates expression of genes involved in AM symbiosis. (**a)** Comparison between phosphate starvation-induced genes (PSI-UP) in WT and genes upregulated in *pho2* mutants in cluster 4 or 6 (*pho2*-UP). (**b)** Comparison of *pho2*-UP with DEGs identified in *PHR2 overexpression* line grown in high phosphate (HP) and PHR2-target genes identified through Chip-Seq ^20^. (**c)** Comparison of *pho2*-UP with core genes known to be important in AM symbiosis ^20^. (**d)** Fold change of genes involved in the strigolactone biosynthesis in uninoculated *pho2* roots relative to WT at 25 (LP) or 250 µM (HP) phosphate treatment. (**e, f)** Heatmap showing the relative expression of Common Symbiosis Signalling Pathway (CSSP, **e**) and phosphate transporter genes (e) in AM-inoculated (+AM) and uninoculated (-AM) plants. The scale is log2-fold change (FC), with red indicating upregulation and blue indicating downregulation. Stars indicate significant changes in the pair-wise comparison (*: p-value ≤ 0.05; **: p-value ≤ 0.01; ***: p-value ≤0.001).

It has previously been shown that PHR2 suppresses PHO2 by activating *miR399* for post-transcriptional silencing ^23, 28, 42, 44, 46, 47^. To investigate whether transcriptional signatures observed in *pho2* mutants resembled those caused by the PHR2-mediated downregulation of *PHO2*, we compared the *pho2-*UP DEGs to those identified to be upregulated in *PHR2* overexpression lines in HP conditions ^20^. While we found genes that were similarly regulated in both *pho2* mutants and by PHR2, 516 genes were distinctly regulated in *pho2* mutants (**Fig.4b**), suggesting that PHO2 activity is not entirely dependent on PHR2-mediated regulation.

Furthermore, comparison of *pho2-*UP to genes identified to be involved in AM symbiosis ^20^ revealed 29 core AM genes that are upregulated in *pho2* mutants in HP (**Fig.4c**). Among these, *pho2* plants had significantly increased expression of genes involved in key stages of SL biosynthesis in HP conditions (**Fig.4d**). Furthermore, CSSP components and a subset of LysM-containing receptor kinases/proteins that are known to be critical for myc factor perception were significantly upregulated in *pho2* mutants (**Fig.4e)**. This suggests that PHO2 suppresses expression of genes involved in pre-symbiotic communication with AM fungi and the subsequent fungal entry into roots.

PHO2 is known to negatively regulate phosphate transporter activity ^45, 46, 47, 48, 49^. Examining the expression of high affinity phosphate transporters (PHT1s), we found that many phosphate transporters involved in both direct and mycorrhizal Pi uptake were significantly upregulated in HP conditions in *pho2* mutants compared to WT (**Fig.4f**), further emphasising the persistence of the phosphate starvation response in *pho2* mutants in HP conditions. The AM-specific phosphate transporter gene, *PT11*, was significantly upregulated in AM-inoculated *pho2* plants in HP, indicating that both direct and mycorrhizal Pi uptake pathways undergo regulation by PHO2.

### PHO2 also suppresses AM symbiosis in the dicot, *Nicotiana benthamiana*

The role of PHO2 in the PSR is largely conserved across monocots and dicots ^23, 42, 43, 44, 47, 48, 52, 53^. To explore whether the *pho2* mutant AM colonisation phenotype is similarly conserved in a dicot, we generated a *pho2* mutant in *N. benthamiana* using CRISPR-Cas9 which targeted the two copies of the *PHO2* gene, *PHO2a* and *PHO2b* (**Fig.S7**). WT *N. benthamiana* plants had reduced colonisation and arbuscule presence compared to *pho2* in LP (25 µM) conditions (**Fig.5a, b**). WT colonisation was further suppressed in HP (1 mM) conditions, whereas *pho2* mutants maintained a similar level of colonisation as in LP. This indicates that the role of PHO2 in suppressing AM colonisation is conserved between rice, a monocot, and *N. benthamiana,* a dicot.

**Figure 5.**
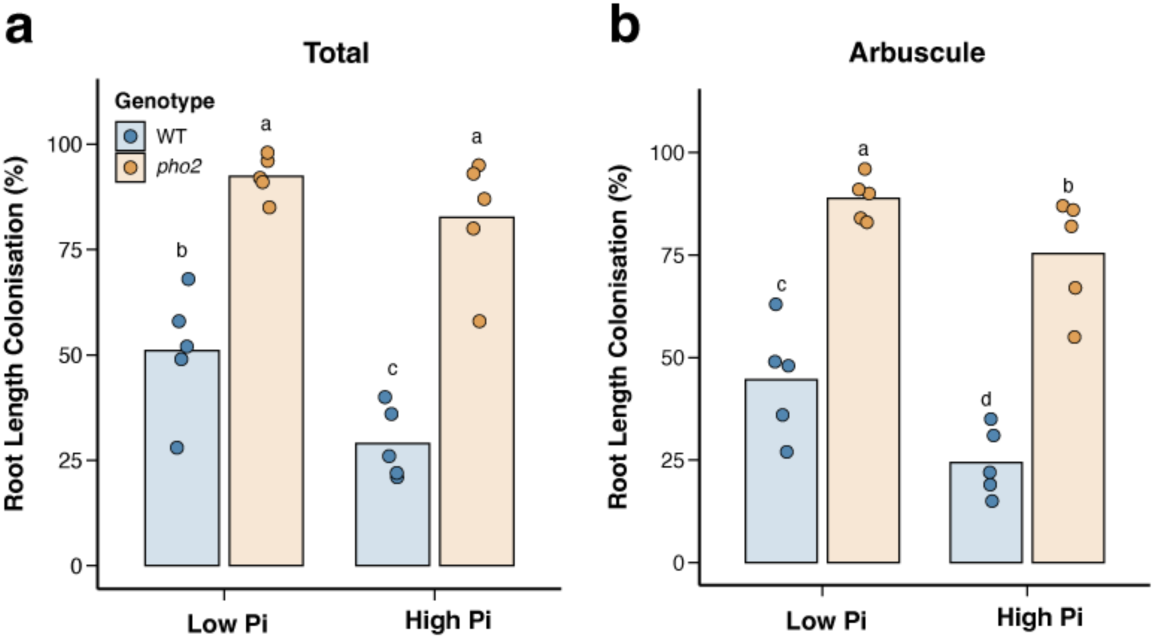
AM colonisation levels of pho2 mutant in *Nicotiana benthamiana*. **(a)** Root length colonisation with of total and **(b)** arbuscules in WT and pho2 *N. benthamiana* at 8 weeks following inoculation with 10 % (w/w) R. irregularis live inoculum. Plants were nourished with half-strength Hoagland’s solution with 25 (Low Pi) or 1000 (High Pi) µM phosphate. Bars show mean values while points indicate individual values. A Kruskal-Wallis test was performed (**(a)** p-value = 0.00154, n = 5; **(b)** p-value = 0.00112, n = 5) and post-hoc pairwise testing using RStudio’s Agricolae package generated letter values showing significantly different groups (p < 0.05).

## Discussion

Recent studies have started to uncover the direct regulation of AM symbiosis by the highly conserved PSR signalling pathway. In Pi-depleted conditions, *PHR* TFs directly activate expression of genes involved in AM symbiosis, whereas their activation is suppressed by SPX proteins in Pi-replete conditions. ^20, 21, 22^. PHO2 has previously been shown to be a key negative regulator of the phosphate starvation response, targeting many phosphate starvation-induced responses, including phosphate transporters. In this study, we reveal that PHO2’s suppression of the PSR also extends to AM establishment in Pi-replete conditions, adding another layer of regulation for Pi-dependent AM inhibition. Crucially, we show that *pho2* mutants in rice exhibit reduced sensitivity to phosphate treatment at a global transcriptome scale (**Fig.3, 4**), accounting for the persistence of phosphate starvation responses and AM genes in high Pi conditions (**Fig.1a, 4a**). To be specific, PHO2 perturbs the expression of genes involved in pre-symbiotic chemical dialogue between plants and the AM fungi and over-accumulates Pi in leaves during AM symbiosis.

Considering the considerable impact of the PSR response in *pho2* mutants, the coordination of PHR2-SPX and PHO2 activities are crucial for optimal AM establishment in varying Pi levels. Non-mycorrhizal studies have shown that *PHO2* is suppressed transcriptionally downstream of PHR2 via miR399 ^23, 42, 44^. The reduced AM colonisation levels in *phr2* ^20^ could be partially due to the elevated *PHO2* transcript levels as *phr2* cannot produce miR399s ^23, 28^. This is consistent with reduced AM colonisation levels in *PHO2* OE lines reported in this study (**Fig.1e**). Conversely, *pho2* mutants might phenocopy *PHR2* OE lines as both lines have displayed higher AM colonisation levels than WT in LP conditions ^20^. The counteraction of miR399 by *IPS1* in LP conditions may account for the more subtle phenotype of symbiotic suppression by PHO2 in LP conditions (**Fig.1b, S3**). Indeed, we observed that the *pho2* mutant displays earlier AM colonisation than WT with occasionally elevated arbuscule abundance (**Fig.1b, S3**). On the other hand, in HP conditions, we consistently observed a strong *pho2* mutant phenotype, retaining high AM colonisation, comparable to WT in LP conditions, highlighting the pivotal role of PHO2 in HP conditions. Interestingly, this phenotype was weaker in unusually HP conditions ([Pi] = 1500 µM) (**Fig.1a, S3, S5a, S6a**), suggesting a threshold for the suppressive activity of PHO2.

We also found that *pho2* mutants in HP conditions form arbuscules that resemble those of WT plants grown in LP conditions in their abundance, size and morphology (**Fig.1b, c, d**, respectively). This contrasts with *PHR2* OE lines which exhibit limited ability to form arbuscules in HP conditions ^20^, suggesting that PHO2 activity is not entirely dependent on regulation by PHR2. Additional evidence for this comes from the portion of PHO2-regulated genes that are distinct from those regulated by PHR2 in AM symbiosis (**Fig.4b**) supporting the assertion that PHO2 contributes further suppression of AM symbiosis independent of PHR2.

PHR2-SPX pathways promote AM colonisation in both early and late stages of AM symbiosis. They control strigolactone production, D14L signalling pathways and the CSSP, which are crucial to pre-symbiotic dialogue between plants and fungi ^9, 10, 11, 12^ and early establishment of infection by AM fungi ^17^. Here we focused on PHO2’s role in the early stage of AM colonisation as the time-course experiment showed that *pho2* mutants are colonised earlier than WT (**Fig.S3**). Analysis of the core set of PHO2-regulated AM genes in the *pho2* root transcriptome revealed that genes involved in strigolactone biosynthesis and the CSSP are elevated in high Pi conditions, indicating host competency to engage with AM fungi (**Fig.4d, e**). This reveals a potential mechanism of PHO2-mediated suppression of AM symbiosis via suppression of pre-symbiotic communication and initial fungal entry into plant roots. As symbiosis was not completely collapsed in WT plants in high Pi, PHO2 likely does not completely abolish their activity, but instead reduces plant permissibility to AM symbiosis. This may reflect the mode of function of PHO2, which targets proteins for degradation, likely in a concentration dependent manner.

Downstream of symbiotic establishment, arbuscule development was also impaired in WT plants in HP conditions but not in the *pho2* mutants (**Fig.1c, d**), implying that elevated PSR in *pho2* enabled arbuscule development or turnover. Considering Pi concentrations in roots were similar in WT and *pho2* mutants (**Fig.S2d**), arbuscule collapse in WT is likely due to the PSR signalling controlled by PHO2, rather than a direct effect of phosphate. However, this requires higher resolution cell-type specific Pi measurements, especially in arbuscule-containing cells.

Finally, we examined the Pi transport activity of *pho2* arbuscules in HP conditions using ^33^P isotope tracing techniques. Considering PHO2 is controlling the phosphate transport between the roots and shoots by targeting PHO1 ^47^, *pho2* mutants enable examination of long-distance transport routes of symbiotically acquired Pi. Our data showed that *pho2* mutants accumulated higher Pi concentrations and total P levels than WT without AM inoculation and this increases further upon AM colonisation (**Fig.2b, S6d**). Furthermore, our ^33^P isotope tracing experiments showed higher concentrations of AM fungal-acquired ^33^P in *pho2* shoots (**Fig.2c**). Taken together, this indicates that arbuscules in *pho2* mutants grown in HP conditions are functional and that PHO2 mediates symbiotic Pi accumulation in the shoot. To identify whether elevated shoot Pi concentrations in *pho2* plants were due to increased Pi uptake in roots, we examined the gene expression of phosphate transporters from our RNA-seq transcriptome. In non-mycorrhizal studies, *pho2* mutants are shown to hyperaccumulate Pi in their shoots ^23, 42, 43, 44^, as is corroborated in our study (**Fig.S2c**). Our data suggests that Pi hyperaccumulation in the leaves of *pho2* plants can be explained by the upregulation of key phosphate transporters involved in both direct and symbiotic Pi uptake pathway (**Fig.4f**). Together with the AM-acquired ^33^P and total phosphorus concentrations of plant shoots, this suggests that *pho2* can simultaneously perform direct and symbiotic Pi uptake. Furthermore, PHO2 likely mediates symbiotic Pi translocation from the root to the shoot in a similar way seen in the direct Pi uptake pathway.

*PHO2* is a conserved gene regulating PSR in monocot and dicot species **(Fig.S1).** We show that *pho2* mutants retain high levels of colonisation under HP conditions in the monocot, rice **(Fig.1a**), and the dicot species, *Nicotiana benthamiana* (**Fig.5a**). This is contradictory to a recent study in the dicot species *Solanum lycopersicum* (tomato) showing the opposite role of PHO2 promoting AM colonisation ^55^. Considering both *N. benthamiana* and tomato are closely related Solanaceae species (**Fig.S1**), this requires further investigation on why and how PHO2 functions in an opposite manner in these species and how prevalent their roles are in other plant species.

In summary, we have identified PHO2 as a key component in the suppression of AM symbiosis in phosphate-sufficient conditions. We demonstrate that loss of PHO2 activity in *pho2* mutants confers reduced sensitivity to external Pi availability, particularly regulating in the early establishment of AM symbiosis, and the accumulation of symbiotically-acquired Pi in the shoot. These findings provide new insights into how plants fine-tune nutrient acquisition strategies and may inform approaches to optimise Pi use efficiency and reduce fertiliser dependence in agriculture.

## Materials and Methods

### Plant materials

All rice lines used in this study are *Oryza sativa L. ssp. Japonica* varieties. The Tos17-retrotransposon insertion lines, *pho2-1* (NF2586) and *pho2-2* (NE9017), were provided by the Rice Genome Resource Centre, Japan. *PHO2* overexpression lines were generated in the *pho2-1* background using the method below. All lines were confirmed as homozygous using PCR-based genotyping (See **Table S2** for primer information).

### Rice growth

Seeds were briefly cleaned in 70 % (v/v with water) ethanol, before rinsing with sterile deionised H2O. Subsequent sterilisation was performed by incubating seeds on a shaker in 3 % sodium hypochlorite (v/v with water) at room temperature for 20 minutes. In a laminar flow hood, sterilised seeds were washed three times in sterile MilliQ water. Seeds were then transferred to 0.6 % bacto-agar (w/v with water) plates and allowed to germinate for 3-5 days at 30 °C in the dark.

Unless stated otherwise, seedlings were transferred to cones containing autoclaved quartz sand. Seedlings received a mock inoculation of RO water or an inoculation of 300 *Rhizophagus irregularis* spores (unless stated otherwise); spores were freshly extracted from hairy carrot (*Daucus carota L.*) root cultures and resuspended in water.

Seedlings were grown in a walk-in controlled environment growth chamber (CONVIRON EVO series) with 12/12 h day/night length and 350 µEm^−2^ light intensity at 28/20 °C with 65 % relative humidity.

For the first week following their transfer to sand, seedlings were watered 3 times weekly with RO water. For subsequent weeks, plants were watered once and fertilised twice weekly with half-strength Hoagland’s nutrient solution containing low (25 µM; LP) or high (250 µM; HP) phosphate unless stated otherwise (**Table S1**). Hoagland’s solution was administered with 0.01 % (w/v) iron supplement, Sequestrene Rapid (Syngenta).

At the point of harvest, roots were washed to remove sand in RO water. Photos were taken and fresh weight of roots and shoots was measured. Samples were taken for fungal quantification and analyses.

To investigate the effects of Pi concentration on fungal colonisation of WT and *pho2* mutant plants, plants received half-strength Hoagland’s containing 25, 250, or 1500 µM inorganic phosphate (**Table S1**). Plants were harvested at 6 weeks post inoculation.

To study AM colonisation of WT and *pho2* mutant plants over a time-course, plants were harvested at 4-, 5-, 6-, and 7-weeks post inoculation.

### AM fungal structure quantification

Representative samples of harvested roots were collected in 10 % KOH (w/v in water), stained with 0.1 % Trypan blue (0.1 % (w/v) Trypan blue, 50 % (v/v) lactic acid, 50 % (v/v) glycerol) and mounted on a glass slide. Roots were observed using brightfield microscopy (LEICA DM750, GT Vision Ltd.) at 200x total magnification and percentage root length colonisation was calculated following a modified gridline intersection method ^56^, counting presence of intraradical hyphae, hyphopodia, arbuscules and vesicles. Total root length colonisation quantified as percentage of root length containing any AM colonisation structure inside the roots.

To capture detailed arbuscule images, harvested root samples were stored in 50 % ethanol (v/v with water) at room temperature before staining with wheat germ agglutinin conjugated to Alexafluor 488 (WGA-Alexafluor 488; 0.2 ug/mL in 1x phosphate-buffered saline solution (PBS; pH7.4)). Stained samples were stored in the dark at 4 °C for at least 6 hours. Immediately prior to imaging, roots were counterstained in propidium iodide (5 ug/mL in 1x PBS) for 1-2 minutes to visualise cell walls. Root fragments were mounted on a glass slide in 1x PBS. Samples were imaged using a Stellaris 8 Falcon confocal microscope (Leica Microsystems GmbH) with excitation by a white light laser at 5 % power at 488 nm. Emission was detected by hybrid detectors at 500-550 nm and 600-700 nm to detect WGA-Alexafluor 488 and propidium iodide emission, respectively. Representative images of arbuscules were obtained using a 40x water immersion objective. To determine arbuscule size, images were captured with a 20x dry objective.

All image processing and analysis was performed using Fiji/ImageJ ^57, 58^. Arbuscule size was determined by dividing arbuscule area by the area of the surrounding cell.

### Quantification of inorganic phosphate

Plants were grown in pots and inoculated with 600 spores of *R. irregularis* or a mock inoculation of RO water. They received half-strength Hoagland’s solution containing 25 µM or 250 µM phosphate (**Table S1**). Plants were harvested at 8 weeks post inoculation, samples were taken for fungal structure quantification and tissue was dried at 50 °C.

18-25 mg of dried tissue was placed in a 2.5 mL tube with two 3mm metal beads and ground to powder (HG-600 Geno/Grinder 2010, SPEX SamplePrep). Inorganic phosphate was extracted by incubation in 500 µL MilliQ water for 45 minutes at 85 °C with occasional inversion.

600 µL of phosphate extraction buffer (0.44 % (w/v) ammonium (para) molybdate, 2.66 % (v/v in RO water) H_2_SO_4_ (36N)) and 100 µL 10 % (w/v) ascorbic acid were added to each sample, before a 1-hour incubation at 42 °C, resulting in a blue colour. Absorbance was measured at 650 nm and phosphate concentration was calculated according to a standard curve of known phosphate concentrations.

### Plant growth conditions for radioisotope tracing of phosphorus

WT, *pho2-1* and *pho2-2* rice seeds were sterilised and germinated as described above. Seedlings were transferred to 1 L pots filled with quartz sand and inoculated with 1200 *Rhizophagus irregularis* spores. Each pot also contained two windowed cores filled with sand ^59^. The windows of the cores were covered with 35 μm nylon mesh (Plastok, UK), which allows fungal hyphal growth into the core interior but excludes root growth ^51^. Following addition of radioisotope described below, one core was rotated to sever AM fungal hyphae entering the core interior. This acted as an internal control for relative contribution of AM symbiosis compared to passive diffusion and alternative microbial cycling processes to plant phosphorus uptake^59^. Plants were grown within a reach-in controlled environment growth chamber (CONVIRON PGR15) with 12/12h day/night length at 28/20°C with 65 % relative humidity. Plants were watered with 25 µM or 250 µM half-strength Hoagland’s, as described above.

### Radioisotope tracing of phosphate transfer from AM fungi to plant

At 8 weeks post-inoculation, 100 µL solution containing 1 Mbq 33P-orthophosphate (111 TBq mmol−1 specific activity, [^33^P]-phosphoric acid (orthophosphate) (^33^P; Hartmann Analytic, Braunschweig, Germany) was added to one of the cores in each pot. 100 µL distilled water was added to the other core. In half the pots in each Pi condition, one core was rotated every 48 hours, while in the other half the cores were kept static. After 2 weeks, plants were harvested. Root tissue samples were collected in 10 % KOH for determining AM colonisation using acidified ink staining, as described below. The remaining shoot and root tissue was separated and freeze-dried for 24 hours (Labogene ScanVac CoolSafe; SLS), weighed, then ground.

### Quantification of ^33^P and total P

Around 30 mg of ground shoot tissue was taken in duplicate and digested using 1 mL of concentrated sulfuric acid overnight. Samples were then heated to 350 °C for 15 minutes, after which they were cooled. 100 µL of hydrogen peroxide was added to the samples and they were heated again for 1 min to decompose residual organic material. Samples were diluted to 10 mL using dH_2_O. A 2 mL aliquot was added to 10 mL of liquid scintillant (Emulsify-safe, PekinElmer). Radioactivity was quantified using liquid scintillation counting (TriCarb® 4900TR, PerkinElmer, Beaconsfield, UK) and ^33^P was calculated using the following equation ^60^ (M^33^ P: mass of ^33^P (mg); cDPM: counts as disintegrations per minute; SAct: specific activity of the source (Bqmmol^−1^); Df: dilution factor; Mwt: molecular mass of P).

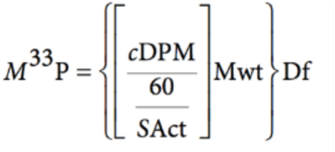

To control for ^33^P transfer via diffusion through the core mesh, the mean ^33^P in plant tissues where the core was rotated was subtracted in each treatment from each ^33^P value in shoots of plants grown in pots with the solution added to static cores ^59^.

To measure the total P content in shoot tissue, 0.25 mL of digest sample was added to a cuvette containing 0.25 mL ammonium molybdate and 0.1 mL of 0.1 M ascorbic acid solutions. This was made up to 1.9 mL with dH_2_O. To make a standard curve, cuvettes with known amounts of P were made up using a 10 mg mL^−1^ P. All solutions were incubated at room temperature in the dark for 45 min before measuring the optical density at 822 nm using a spectrophotometer (Jenway 6300).

### Quantification of AM colonisation from ^33^P tracing experiment

Root samples were heated in around 1 mL of 10 % (w/v) KOH at 90 °C for 30 min before being rinsed in distilled water to remove any residual substrate. Root samples were then incubated in around 1 mL of 0.3 M HCl for between 15 – 120 min. New HCl was added and the incubation was repeated for 15 min to further clarify the root tissue. HCl was removed and around 1 mL of staining solution (50 % (v/v) glycerol, 10 % (v/v) black ink, 2.5 % (v/v) acetic acid) added to the root samples which were then incubated at 90°C for 5 min. The staining solution was removed and destaining solution (50 % glycerol, 2.5 % acetic acid) was added. Samples were incubated at room temperature overnight. Around 1 cm root pieces were mounted in destaining solution on a glass slide. Percentage root length colonisation was quantified as described above.

### RNA Extraction, cDNA synthesis and quantitative Reverse Transcriptase (qRT-)PCR

Approximately 60-100 mg of shoot or root tissue from each sample was collected in 2 mL safe-lock Eppendorfs tubes containing two 3 mm autoclaved metal beads. The tissue was immediately frozen in liquid nitrogen and stored at –80 °C. Frozen samples were ground to powder (HG-600 Geno/Grinder 2010, SPEX SamplePrep) and RNA was extracted using the RNeasy Plant Minikit (QIAgen), following the manufacturer’s instructions. RNA concentration was quantified by spectrophotometry (NanoDrop One C, Thermo-Fisher Scientific, USA). Absence of genomic DNA in extracted RNA was confirmed using PCR and RNA integrity was checked by gel electrophoresis on a 1 % agarose gel at 120 V.

For samples used in cDNA synthesis, genomic DNA contamination was removed using DNase I (1 U/uL, ThermoScientific, USA). cDNA was synthesised using SuperScript IV reverse transcriptase and RNase-OUT™ (Invitrogen, USA). Success of cDNA synthesis and absence of genomic DNA contamination were confirmed using PCR. Quantitative qPCR was performed with primers in **Table S2** using a master mix of GoTaq DNA Polymerase® (Promega) and SYBR™ green (Thermo-Fisher Scientific) with a Bio-Rad CFX384 qPCR cycler (Bio-Rad Laboratories, Shanghai, China).

### RNA-seq and data analysis

RNA extraction and quality control were performed as described above. Genomic DNA contamination was removed from samples for RNA-seq using TURBO DNase™ (Invitrogen). mRNA library preparation and 150 bp paired-end sequencing were performed by Novogene (UK Company Limited, Cambridge). Raw sequencing reads were quality-checked with FastQC (v0.12.1) and subsequently adapter-trimmed using Trimmomatic (v0.39) ^61^. High-quality reads were then quantified at the transcript level using SALMON 0.7.0 ^62^ quasi-transcript mapping. This was performed within the Mambaforge environment (https://mamba.readthedocs.io/en/latest/installation/mamba-installation.html) on Windows. Reads were mapped to the *Oryza sativa* MSU v7.0 reference transcriptome downloaded from Phytosome v.12 (https://phytozome-next.jgi.doe.gov/). Transcript-level abundance estimates from Salmon were imported into R (v4.3.1) and summarised to the gene level using the tximport package (v1.28.0) in R/Bioconductor 3.21 (https://www.bioconductor.org/packages/release/bioc/html/tximport.html) before input into DESeq2 (v1.40.0) ^63^ in R/Bioconductor (https://bioconductor.org/packages/devel/bioc/vignettes/DESeq2/inst/doc/DESeq2.ht ml). DESeq2 was used for differential expression analysis with cut-offs for differentially expressed genes (DEGs): FDR ≤ 0.05 and an absolute log_2_FoldChange ≥ log_2_(1.5). DEGs were grouped using the k-means clustering. The resulting clusters were visualised as heatmaps generated using ComplexHeatmap (v2.16.0) ^64^ in R/Bioconductor 3.22 (https://www.bioconductor.org/packages/devel/bioc/html/ComplexHeatmap.html). Gene ontology (GO) enrichment of gene clusters was done using AgriGO ^65^. AM-related genes were identified using a published gene list ^20^.

### Genotyping

Leaf tissue was collected in 2 mL Eppendorfs tubes with three 3 mm autoclaved glass beads and flash frozen in liquid nitrogen. Tissue was ground to powder (HG-600 Geno/Grinder 2010, SPEX SamplePrep). DNA extraction was performed by incubating at 98 °C for 10 minutes with a simple DNA extraction buffer (1 M KCl, 0.1 M Tris, 10 mM EDTA, adjusted to pH 7.4 with HCl). Samples were vortexed briefly then centrifuged for 5 minutes at 13000 rpm and supernatant was transferred to 1.5 mL tubes with 100 % isopropanol and inverted. Samples were then centrifuged for 15 minutes (13000 rpm) to form pellets. The pellet was washed twice with 70 % (v/v) ethanol and air-dried at room temperature before being resuspended in 30 mL autoclaved RO water. Genotyping was performed by PCR with primers listed in **Table S2**.

### Generating pho2 mutant in Nicotiana benthamiana

*Arabidopsis thaliana PHO2* (AT2G33770) was used to query against the *Nicotiana benthamiana* genome v1.0.1 ^66^ using the tBLASTn algorithm v2.10.1 + ^67^. Two genes were identified with high sequence similarity (Nb*PHO2a* and *b*) and supported by the phylogenetic tree (**Fig.S1**). One gRNA was designed to target both *PHO2* sequences and mutants were generated as previously described ^68^. Briefly, Tobacco Rattle Virus vectors were assembled expressing the *NbPHO2* gRNA augmented with a mobile motif and used to infect *N. benthamiana* plants expressing SpCAS9. After approximately 2 months, seeds were collected, and gene editing was confirmed by the Sanger sequencing of a PCR product at the gRNA target location (see primers in **Table S2**). One plant was identified with homozygous one base pair insertion in each of the two homologs, resulting in premature stop codon (**Fig.S7**). The specific target sequences of the guide RNAs, and the mutations present in each mutant plant, are listed in **Table S3**.

### AM colonisation assay of *Nicotiana benthamiana pho2* mutant plants

WT and *Nbpho2a/b* double mutant plants were assessed for engagement with AM fungi by placing seeds into a 40 mL cone system containing 10 % (w/w) live inoculum. Live inoculum was generated by inoculating WT plants with 5 % (w/w) crude inoculum (produced on *Tagetes multiflora*) of *R. irregularis* and growth in 4 x 4 x 4.5 cm^3^ pots. Plants were grown at 22 °C/20 °C,12 h light/12 h dark cycle with 60 % relative humidity, nourished with a half-strength Hoagland’s solution containing 25 μM Pi twice a week. After approximately 2 months, live inoculum sand and root mixture was produced by removing all live shoots and chopping the existing root system. Cones containing wild-type or *Nbpho2a/b* plants were grown in the same growth condition described above and subjected to a twice a week nutrient treatment of either 25 μM or 1 mM Pi half-strength Hoagland’s solution for 6 weeks. The inoculated roots were harvested and stained with Trypan blue dye for visualising fungal structures. Due to small root systems, two plants were combined as one biological sample and mounted on slides to quantify colonisation levels according to the modified gridline intersection method ^56^.

### Generating *PHO2*-OE lines in rice

An Os*PHO2* expression cassette was assembled by Goldengate cloning ^69^, transferred by Gateway recombination into binary vector pRLF12-R1R2 to create pEW583, followed by transfer to *Agrobacterium tumefaciens* strain EHA105 as previously described ^70^. Callus for transformation of the *pho2-1* mutant was generated by plating surface-sterilised mature seed, with embryo axes removed, on N6DT medium (3.95 g/L N6 basal salts, 30g/L sucrose, 300mg/L casein hydrolysate, 100mg/L myo-inositol, 2878mg/L proline, 0.5mg/L nicotinic acid, 0.5mg/L pyridoxine HCl, 1mg/L thiamine HCl, 37.3 mg/L Na2EDTA, 27.8 mg/L FeSO4, 2mg/L 2,4-D Na salt, 150mg/L Timentin, 4g/L Gelrite, pH5.8). Plates were sealed with Parafilm and cultured in the dark at 28°C for 21 days, after which time callus was cut into 2-4mm pieces, plated on fresh N6DT and cultured as before for a further 4 days. Transformation of the rice callus material was carried out as previously described ^70^.

## Author Contributions

Conceptualisation, S.B., J.C.; Investigation, S.B., S.P., J.C. E.E.; Methodology, S.B., S.P., E.E., N.F., N.M., A. W., M.B-W., T.R., B.P., M.W., M.S.H., E.W., K.F.; Writing, S.B., S.P., J.C.; Funding acquisition, E.E.E., J.C.

All authors contributed and approved the current manuscript.

## Acknowledgments

We thank Aliya Santosa, Shawna Rowe, Abigail Brock, Chongjing Xia, research support staff of Crop Science Centre, and Ruth Bates (Niab) for their technical support.

S.B. was supported by the Frank Smart - School of Biological Science PhD Scholarship. J.C., S.P. and N.M. and rice transformation activity at Niab were supported by the Royal Society University Research Fellowship (URF\R1\21134). E.E.E. was supported by NSF Postdoctoral Research Fellowship in Biology Grant No. 2305688. T.R. was supported by the summer internship programme of the Department of Plant Sciences, University of Cambridge. K.F. and A.W. are supported by the ERC CoG “MYCOREV” (865225), M.B-W. is supported by a University of Sheffield PhD studentship. We thank the de Laszlo Foundation for generously supporting PhD student research at the University of Sheffield.

