## Supplementary files for "PHOSPHATE OVERACCUMULATOR 2 (PHO2) is a negative regulator of arbuscular mycorrhizal symbiosis"

Birch et al.

### Supplementary Materials

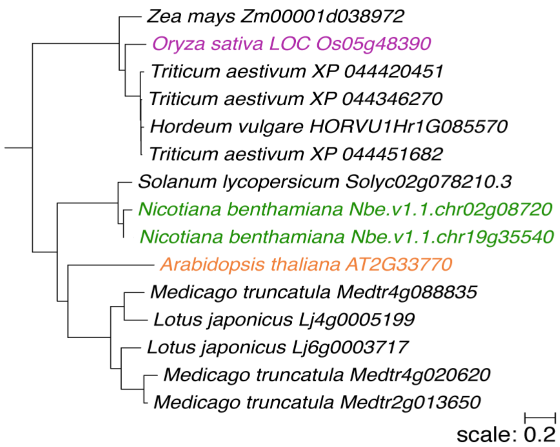


**Figure S1. A phylogenetic tree of PHO2 proteins in 9 plant species:** Arabidopsis (*Arabidopsis thaliana*, orange colour), Medicago (*Medicago truncatula*), Lotus (*Lotus japonicus*), tobacco (*Nicotiana benthamiana*, green), tomato (*Solanum lycopersicum*), rice (*Oryza sativa*, purple), barley (*Hordeum vulgare*), wheat (*Triticum aestivum*), and maize (*Zea mays*).

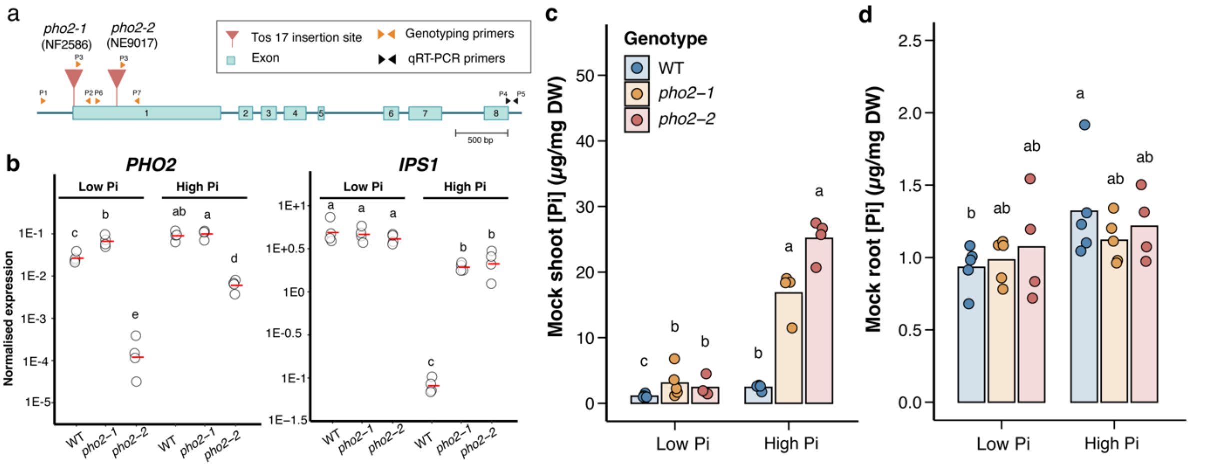


**Figure S2. Allelic characterisation of rice *pho2* mutants.**

**(a)** Position of Tos17 insertion sites in *pho2* mutants. (**b)** Expression of *PHO2* and phosphate starvation response marker gene, *IPS1,* in WT, *pho2-1* and *pho2-2* plants, normalised to the expression of 3 housekeeping genes: *OsCYCLOPHILIN2, OsGAPDH* and *OsPOLYUBIQUITIN*. Plants were grown in cones for 4 weeks and received half-strength Hoagland's solution containing 25 (Low Pi) or 250 (High Pi) µM phosphate. Points indicate individual samples and red lines show the mean for each genotype. **(c)** Shoot and (**d)** Root inorganic phosphate (Pi) concentrations of uninoculated (plants grown in pots for 8 weeks and fertilised with half-strength Hoagland's containing 25 (Low P) or 250 µM (High P) phosphate. Bars indicate the mean value for each genotype, and points show individual biological replicate. Groups were compared using Kruskal-Wallis tests (**(b)** *PHO2:* p-value = 3.63E-08; *IPS1:* p-value =3.05E-08; **c)** p-value = 2.67E-05; **d)** p-value = 0.0205 and n = 4 or 5) and post-hoc pairwise comparison using RStudio's Agricolae package generated letter labels denoting significantly different groups ($p < 0.05$).

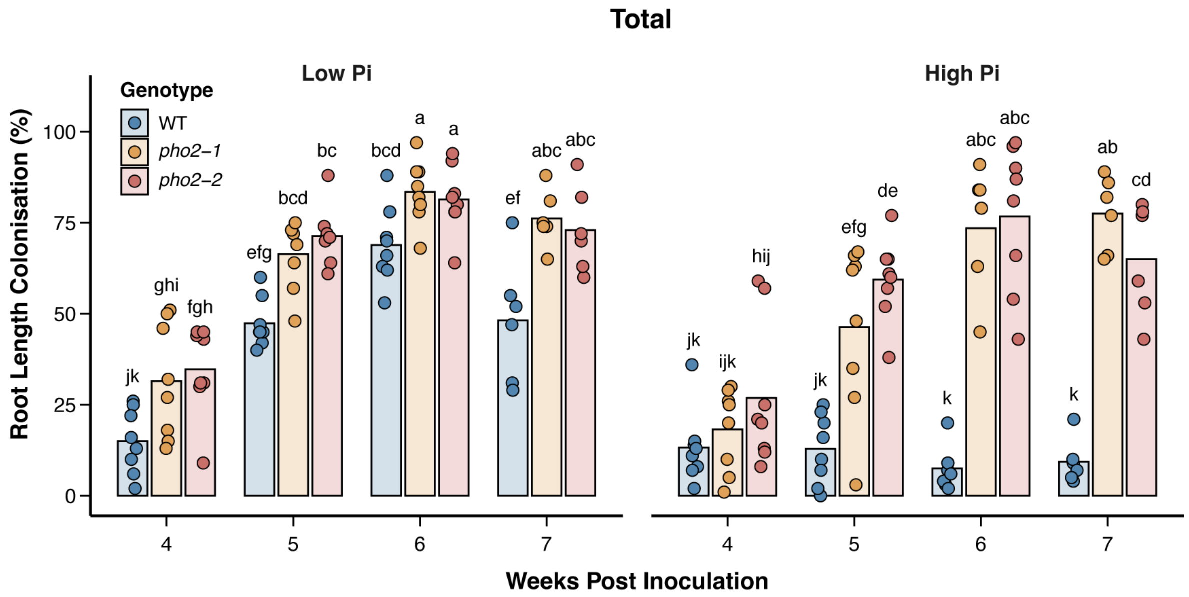


**Figure S3. Time-course AM colonisation levels of *pho2* mutants.**

Total root length colonisation at 4, 5, 6 and 7 weeks post inoculation (wpi) in WT and *pho2* mutant rice plants inoculated with 300 spores of *Rhizophagus irregularis* and fertilised with half-strength Hoagland's solution containing 25 (Low Pi) or 250 (High Pi) µM phosphate. Bars indicate the mean for each genotype at each timepoint, and points show individual biological replicates. Kruskal-Wallis tests were performed across Low and High Pi samples (p-value = 0, n = 8). Letters indicate significantly different groups ($p < 0.05$) based on post-hoc pairwise comparison using the Agricolae package in RStudio.

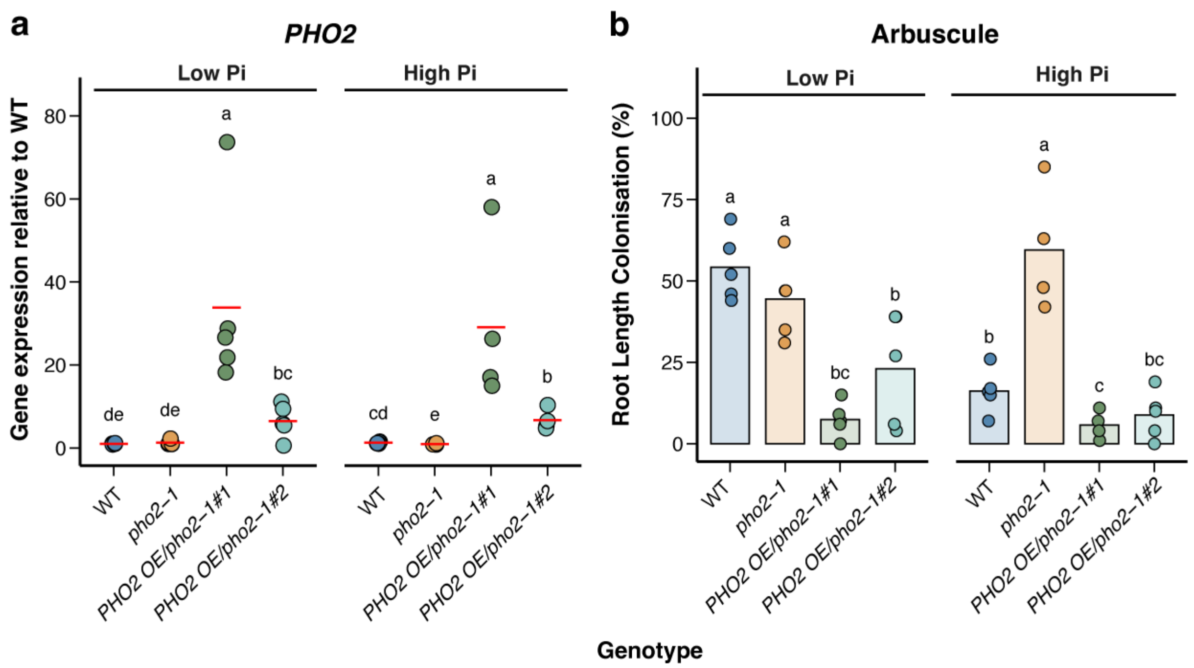


**Figure S4.** ***PHO2*** **expression and AM colonisation of** ***PHO2*****-overexpressing lines.**

**(a)** Expression of *PHO2*. Expression of *PHO2* was normalised to the expression of 3 housekeeping genes: *OsCYCLOPHILIN2, OsGAPDH and OsPOLYUBIQUITIN,* and divided by the mean value for WT at Low Pi. Points indicate individual samples, and red lines show the mean for each genotype. *(***b)** Root length colonisation levels of arbuscules PHO2 overexpression lines in a *pho2-1* mutant background under the maize ubiquitin promoter. Plants were grown in cones for 6 weeks following inoculation with 300 spores of *R. irregularis* and received half-strength Hoagland's solution containing 25 (Low Pi) or 250 µM (High Pi) phosphate. Bars indicate the mean value for each group and points show individual samples. For statistical analysis, groups were compared using a Kruskal-Wallis test (**(a)** p-value = 0.000209 and (**b)** p-value = 0.000233, n = 4 or 5) and post-hoc pairwise comparison using RStudio's Agricolae package generated letter labels denoting significantly different groups ($p < 0.05$).

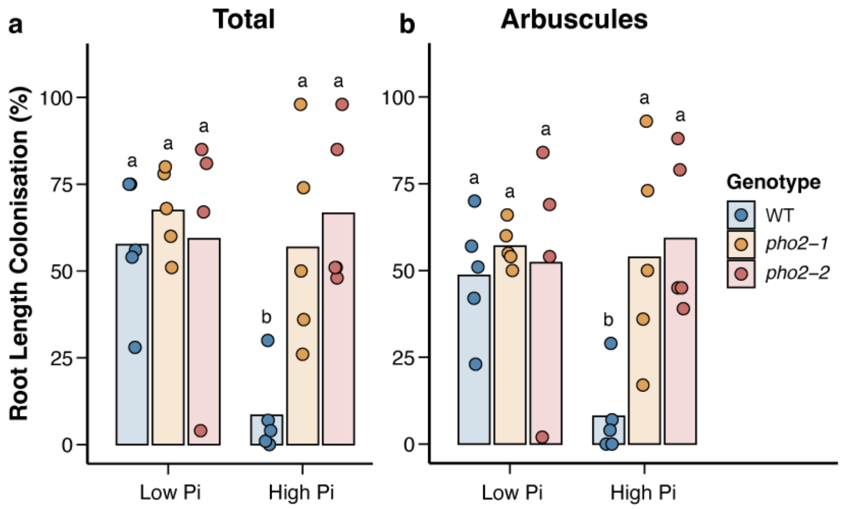


**Figure S5. AM colonisation levels of *pho2* mutants grown for mycorrhizal growth promotion effect.**

**(a)** Root length colonisation levels of total and (**b)** arbuscules of plants grown for 8 weeks in 400 mL pots, receiving half-strength Hoagland's solution containing 25 (Low Pi) or 250 μM (High Pi) phosphate. Bars show mean values while points indicate individual values. A Kruskal-Wallis test was performed(**(a)** p-value = 0.00341 and (**b)** p-value = 0.00573, n = 4 or 5) and post-hoc pairwise testing using RStudio's Agricolae package generated letter values showing significantly different groups (p <0.05).

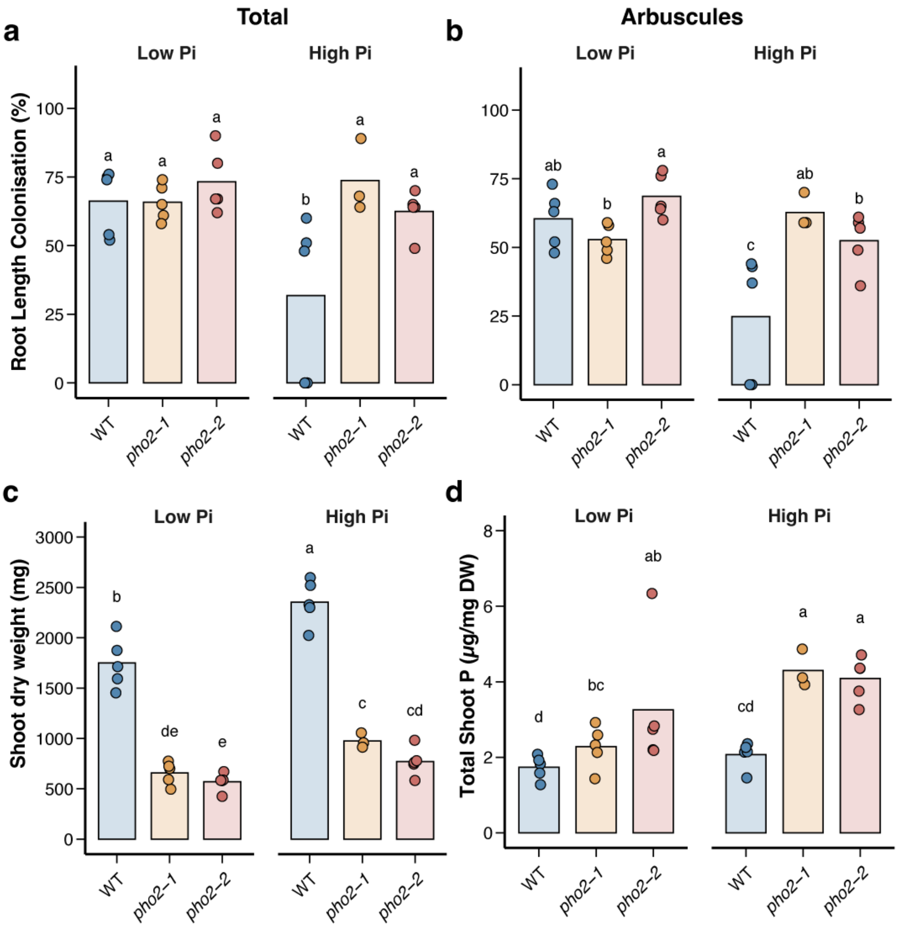


**Figure S6. AM colonisation, shoot dry weight and total shoot phosphorus levels of *pho2* mutants to investigate symbiotic phosphate uptake.**

**(a)** Root length colonisation of total **b)** arbuscules. **(c)** Shoot dry weight **(d)** Total shoot phosphorus concentration of plants in $^{33}$P mycorrhizal phosphate uptake assay. Plants were grown in 1 L pots and received half-strength Hoagland's containing 25 (Low Pi) or 250 µM (High Pi) phosphate. At 8 weeks post-inoculation, $^{33}$P-orthophosphate was administered to an AM-accessible core in each pot, and plants were harvested 2 weeks later. Bars show mean values while points indicate individual values. A Kruskal-Wallis test was performed ((**a)** p-value = 0.0319, (**b)** p-value = 0.00278, (**c)** p-value = 2.58E-08 and (**d)** p-value = 0.00161, n = 3, 4 or 5). Post-hoc pairwise testing using RStudio's Agricolae package generated letter values showing significantly different groups ($p < 0.05$).

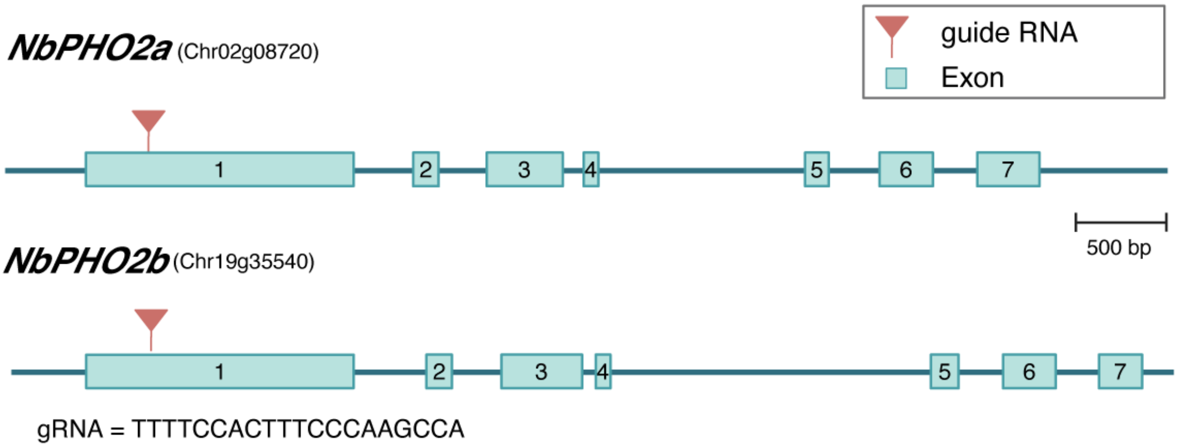


**Figure S7.** ***pho2 CRISPR*** **mutants in** ***Nicotiana benthamiana*****.**

Diagram showing the gene structure of two *PHO2* homologs in *Nicotiana benthamiana*. Each box represents an exon flanked by introns (space between exons). The location of guide RNA (see sequence in **Table S3**) targeting both pho2 homologs is marked with a red triangle. The width of each component is proportional to the to scale (500 bp).

**Table S1. Nutrient concentrations in half-strength Hoagland's solutions used in this study.**

| | | Phosphate Concentration in half-strength Hoagland's solution (μM) | | | |
|---|---|---|---|---|---|
| | | 25 | 250 | 500 | 1500 |
| Macronutrient concentrations (mM) | $KNO_3$ | 2.5 | 2.5 | 2.5 | 1.5 |
| | $Ca(NO_3)_2$*$4H_2O$ | 0.25 | 0.25 | 0.25 | 0.75 |
| | $KH_2PO_4$ | 0.025 | 0.25 | 0.5 | 1.5 |
| | KCl | 0.475 | 0.25 | 0 | 0 |
| | $CaCl_2$ | 0.5 | 0.5 | 0.5 | 0 |
| Micronutrient concentrations (μM) | $MnSO_4$*H2O | 40 | | | |
| | $ZnSO_4$*$7H_2O$ | 3.5 | | | |
| | $MgSO_4$*$7H_2O$ | 1.6 | | | |
| | $Na_2B_4O_7$*$10H_2O$ | 260 | | | |
| | $(NH_4)6Mo_7O_{24}$*$4H_2O$ | 0.4 | | | |

**Table S2. Primers used in this study**

| Purpose | Gene | ID | Primer name | Primer sequence | Reference |
|---|---|---|---|---|---|
| Genotyping of *Ospho2-1* | WT | LOC_Os 05g48390 | P1 | ACACCAGGAAGTCCAGAACG | This study |
| | | | P2 | CCAGAGCTCGTTTCCAAGTC | |
| | *Ospho2-1* | | P3 | AGTCGCTGATTTCTTCACCAAGG | |
| | | | P2 | CCAGAGCTCGTTTCCAAGTC | |
| Genotyping of *Ospho2-2* | WT | | P6 | GACTTGGAAACGAGCTCTGG | |
| | | | P7 | GGCATAAGGGAAGCATGAAA | |
| | *Ospho2-2* | | P3 | AGTCGCTGATTTCTTCACCAAGG | |
| | | | P7 | GGCATAAGGGAAGCATGAAA | |
| Genotyping of *Nbpho2* | *nbpho2a* | Chr02g08 720 | pEEE372 | CGTGTGTAATTTTCAAGTCTTCACTC | |
| | | | pEEE374 | CTGAACTTGAACCCTTTGTCC | |
| | *nbpho2b* | Chr19g35 540 | pEEE383 | GCTCTTGATTTTGAATAATTATAGTTCATT GC | |
| | | | pEEE374 | CTGAACTTGAACCCTTTGTCC | |
| qRT-PCR | *OsPHO2* | LOC_Os 05g48390 | P4 | AAGCTCTTGCCGAAGCTTGT | |
| | | | P5 | TTGGTGTCCCGATTTTGTCAC | |
| | *Ri Ef 1a* | | F | GCTATTTTGATCATTGCCGCC | Perez Tienda et al., 2014 |
| | | | R | TCATTAAAACGTTCTTCCGACC | |
| | *OsGAPDH* | LOC_Os 08g03290 | F | CTGATGATATGGACCTGAGTCTACTTTT | Gutjahr et al., 2008 |
| | | | R | CAACTGCACTGGACGGCTTA | |
| | *OsUbiquitin* | LOC_Os 06g46770 | F | CATGGAGCTGCTGCTGTTCTAG | Gutjahr et al., 2008 |
| | | | R | CAGACAACCATAGCTCCATTGG | |
| | *OsCyclophillin* | LOC_Os 02g02890 | F | GTGGTGTTAGTCTTTTTATGAGTTCGT | Gutjahr et al., 2008 |
| | | | R | ACCAAACCATGGGCGATCT | |
| | *OsIPS1* | LOC_Os0 3g05334 | F | TTGGCAATTATTCGGTGGAT | Yang et al., 2012 |
| | | | R | ACCATTTCACCATCCTCTTTATG | |

Perez-Tienda et al., Plant Physiol Biochem 75 :1-8 (2014)
Gutjahr et al.,  The Plant Cell 20 : 2989-3005 (2008)
Yang et al., The Plant Cell 24: 4236-4251 (2012)

**Table S3. Guide RNAs, resulting mutations and sequence information in *Nicotiana benthamiana pho2* mutants used in this study**

| | Target 1 | Target 1 Result | Target 1 Sequence | Target 2 | Target 2 Result | Target 2 Sequence |
|---|---|---|---|---|---|---|
| gEEE431-1-10-1 | *Nbpho2a* | +T | CCATGGCTTTGG GAAAGTGGAAAA | nbpho2b | +G | CCATGGGCTTGG GAAAGTGGAAAA |

| Sequences | |
|---|---|
| *NbPHO2a* | Chr02g08720 |
| *NbPHO2b* | Chr19g35540 |
| *NbPHO2* gRNA1 | TTTTCCACTTTCCCAAGCCA |
